# Rice lectin protein *Osr40c1* imparts drought tolerance by modulating *Os*SAM2, *Os*SAP8 and chromatin-associated proteins

**DOI:** 10.1101/2020.04.19.049288

**Authors:** Salman Sahid, Chandan Roy, Soumitra Paul, Riddhi Datta

**Author notes:** Email addresses: Salman Sahid, Chandan Roy, Soumitra Paul, Riddhi Datta.

## Abstract

Lectin proteins play an important role in biotic and abiotic stress responses in plants. Although the rice lectin protein, *Os*r40c1, has been reported to be regulated by drought stress, the mechanism of its drought tolerance activity has not been studied so far. In this study, it has been depicted that expression of *Osr40c1* gene correlates with the drought tolerance potential of various rice cultivars. Transgenic rice plants overexpressing *Osr40c1* were significantly more tolerant to drought stress over the wild-type plants. Furthermore, ectopic expression of the *Osr40c1* gene in tobacco yielded a similar result. Interestingly, the protein displayed a nucleo-cytoplasmic localization and was found to interact with a number of drought-responsive proteins like *Os*SAM2, *Os*SAP8, *Os*MNB1B, and *Os*H4. Fascinatingly, silencing of each of these protein partners led to drought susceptibility in the otherwise tolerant *Os*r40c1 expressing transgenic tobacco lines indicating that these partners were crucial for the *Os*r40c1-mediated drought tolerance *in planta*. Together, the present investigation delineated the novel role of *Os*r40c1 protein in imparting drought tolerance by regulating the chromatin proteins, *Os*MNB1B and *Os*H4, which presumably enables *Os*SAP8 to induce downstream gene expression. In addition, its interaction with *Os*SAM2 might induce polyamine biosynthesis thus further improving drought tolerance in plants.

**Highlights:** A rice lectin protein, *Osr*40c1, plays a crucial role in imparting drought stress tolerance in plants by modulating *Os*SAM2 as well as the transcriptional regulators *Os*SAP8, *Os*MNB1B and *Os*H4.

## Introduction

Rice is a staple food crop in most of the countries of Southeast Asia (Cassman *et al*., 2003; Seck *et al*., 2012). Generally, rice is cultivated in irrigated land and requires a higher amount of water compared to other crops (Mohanty *et al*., 2013). Rice cultivation is hampered when the plants are exposed to a period of water deficiency due to an insufficient water supply or uncertainty of rainfall. In addition, plants are always exposed to a plethora of environmental challenges. Among them, drought stress has serious impacts on several physiological functions of plants. It inhibits growth and hampers seed development (Atkinson and Urwin, 2012; You *et al*., 2014). Therefore, the production of large biomass or high grain yield under water deficit or drought stress conditions has always been a major challenge. During drought, plants regulate several metabolic pathways involved with enzyme activity, alteration in different metabolite levels, and accumulation of compatible solutes to withstand the condition. However, the first impression of drought stress in plants is found from their morphological changes.

Drought tolerance is a complex physiological phenomenon that is not controlled by a single protein but involves the interaction of different signaling pathways (Datta et al. 2020). Plants have carbohydrate-binding lectin protein families that play an important role in plant defense and abiotic stress response including drought stress (Li *et al*., 2014). In plants, the lectin family proteins can be classified into seven subfamilies (Peumans *et al*., 2001). However, recently 12 plant lectin families have been recognized (Fouqueart *et al*., 2008). Out of these families, R40 family of lectin protein contains the carbohydrate-binding ricin-like domain and shows osmotic stress responsiveness. In rice, five functional *Os*R40 proteins, r40c1, r40c2, r40g2, r40g3 and putative r40c1 have been reported (Jiang *et al*., 2012). Fascinatingly, the R40 family of proteins has been reported to be up-regulated by abscisic acid (ABA) treatment and to play a salinity responsive role in rice (Moons *et al*., 1995; Moons *et al*., 1997). Subsequently, it has been observed that the *Os*R40 family members exhibit osmotic stress-responsive function and are up-regulated during drought and salt treatment in plants (Riccardi *et al*., 2004; Jiang *et al*., 2012). In addition, it has also been demonstrated that a putative r40c1 protein (LOC_Os03g21040) is up-accumulated in the roots of DREB2A-overexpressing rice plants under drought stress (Paul *et al*., 2015). However, the functional mechanism of the *Os*r40c1 protein in regulating drought stress has not yet been studied. Therefore, the present study aims to unravel the molecular mechanism of this putative *Os*r40c1 protein in regulating drought stress response *in planta*.

In this work, we have reported that the expression of the *Osr40c1* gene is directly correlated with the drought tolerance potential of different rice cultivars and its overexpression significantly enhances drought tolerance in rice. In tobacco, the ectopic expression of the *Osr40c1* has also been found to impart drought tolerance. Moreover, it has been identified that the protein interacts with several drought-responsive protein partners like *OsSAP8, OsMNB1B, OsSAM2*, and *OsH4*. The silencing of these interacting protein partners has been found to induce susceptibility to drought stress thus demonstrating that the partners are indispensable for the *Os*r40c1-mediated drought stress tolerance *in planta*.

## Materials and methods

### Plant material and stress treatment

The eight *indica* rice cultivars, namely IR36, Ranjit, Khitish, IR64, IR72, Swarna sub1, Jaldi13, and MTU1010 were selected and seeds were procured from Chinsurah Rice Research Station, West Bengal, India. Plants were grown in the net-house under optimum condition. Plants were subjected to drought stress at vegetative stage (60 days old) with complete withdrawl of water for 7 days till the moisture level was maintained upto 45 % to 50 % (Paul et al., 2015). After 7 days of drought exposure, plants were re-watered for 24 hrs. In addition, rice cultivars were germinated and grown on filter papers wetted with water for 10 days under optimum photoperiod at 37 °C. The seedlings were treated with 20% PEG (6000) solution for 7 days for osmotic stress treatment.

Tobacco (*Nicotiana tabacum L. cv. Xanthi*) seeds were germinated and grown in standard Murashige and Skoog (MS) medium under 16 h light/ 8 h dark photoperiod as standardized before (Murashige and Skoog, 1962; Ghanta *et al*., 2013). For drought stress, 60 days old soil-grown plants were subjected to drought stress by withdrawal of watering for 5 days until soil moisture content reached 45 %.

### Morphological Analysis

Following the 7 days of drought stress, plant height, number of tillers, number of leaves, percentage of rolled and brown leaves, root length, dry weight of roots and shoots were considered under control, drought and re-watered conditions. Shoots and roots were dried at 50 °C for 3 days to measure the dry weight.

### Biochemical analyses

Proline content was estimated from the roots and shoots of rice plants collected from control, drought, and re-watered condition following Woodrow *et al*. (2016). Briefly, the roots and shoots were homogenized in 80% ethanol and mixed with 1% ninhydrin solution. The optical density was recorded by spectrophotometer (Hitachi) at 595 nanometer. The shoots and roots were homogenized in 0.05 % toluene and glycine betaine was measured according to Grieve and Grattan (1983). The total soluble sugar content was estimated following Dubois,1956 with slight modifications. The samples were homogenized, boiled for 30 minutes and centrifuged (Hitachi) at 5000 rpm for 5 minutes. The supernatant was mixed with 9% phenol and 98% sulfuric acid and optical density has been measured through spectrophotometer (Hitachi) at 485 nanometer. The ascorbic acid and GSH content of roots and shoots from similar set of plants were also analyzed (Gillespie and Ainsworth 2007; Chen et al., 2011).

### RNA Extraction and quantitative RT PCR

Total RNA was extracted from root samples using the Trizol method. Complementary DNA (cDNA) was synthesised using iscript cDNA synthesis Kit following the manufacture’s protocol (Bio-rad). The quantitative PCR (qPCR) analysis was perfomed in CFX 96 Real time PCR (Bio-rad) using gene specific primers (Table S1) and iTaq Universal SYBR Green Supermix (Bio-rad). The actin gene (*OsAct1*) was considered as referrence gene while *OsDehydrin* gene has been selected as drought-responsive marker gene.

### Generation of *Osr40c1* overexpressing transgenic rice plants and stress assay

Total RNA was extracted from rice roots and cDNA was prepared using cDNA synthesis kit (Bio-rad). The full length gene was amplified from cDNA with gene specific primer (Fig S1) to obtain 1047 bp fragment of *Osr40c1* (Genebank Accession no: AC126222.2). The amplified fragment was cloned into pGEMT-Easy vector system (Promega) and then sub-cloned into the *EcoRI* and *BamHI* restriction enzyme (RE) sites of *pEGAD* vector under the control of CaMV35s promoter. The recombinant construct (*35S*::Os*r40c1*-*eGFP*) was introduced into *Agrobacterium* strain *GV3101* for transformation of *indica* rice cultivar khitish (IET4095) following Datta et al. (2000). The *Agrobacterium* strain containing the empty *pEGAD* vector (*35S::eGFP*) was also used for transformation of rice plants to generate vector control (VC) lines. The putative transgenic lines along with wild-type (WT) were grown in greenhouse with optimum photoperiodic conditions (16h light and 8h dark). The transgenic lines were screened by genomic DNA PCR using *Bar* gene specific primer. The positive transgenic plants were maintained upto T2 generation. Different morphological parameters, metabolite contents and expression of *Osr40c1* were analyzed from trangenic, VC and WT plants (grown under control and drought condition) as described above.

### Subcellular Localization of *Os*r40c1 Protein

*Agrobacterium-mediated* infiltration of onion epidermeal cell and subcellular localization of proteins was performed according to Xu et al. (2014). The Agrobacterium strain GV3101 harbouring the recombinant construct (*35S::Osr40c1-eGFP*) was used to infiltrate the onion epidermal cells. The infiltrated onion cells were incubated in dark at 28 °C. The epidermal cells were also stained with the nuclear stain, DAPI and visualized under confocal laser scanning microscope.

### Generation of ectopic line of *Osr40c1* in tobacco

The recombinant construct (*35S::Osr40c1-eGFP*) was used for *Agrobacterium*-mediated transformation of tobacco following leaf disc method (Ghanta *et al*., 2013). The putative transformed lines were confirmed through genomic DNA PCR with *Bar* gene specific primers. The positive transgenic lines were grown up to T_2_ generation. The morphological and biochemical parameters as well as Osr40c1 expression were analyzed in response to stress treatment as described above.

### Yeast Two Hybrid Assay

Yeast two-hybrid analysis was performed using the Matchmaker Gold Yeast Two-Hybrid System (Clontech) following manufacturer’s protocol. The *Osr40c1* gene was cloned into *Eco*RI and *BamHI* RE sites of *pGBKT7* vector to generate recombinant *BD-Osr40c1* bait construct. Yeast transformation was performed according to the instructions of clonetech manual. The cDNA library was prepared from rice roots collected from the plants exposed under drought stress. The cDNA library was ligated to *pGADT7-Rec* vector to prepare recombinant AD construct using Make Your Own “Mate and Plate” Library system (Clontech). The yeast strain Y2H gold was co-transformed with bait and prey recombinant construct and colonies were screened against DDO (SD/-Leu/-Trp) and QDO/X/A (SD/-Ade/-His/-Leu/-Trp with aureobsidin A and X-α-gal). Selected blue colonies were analyzed by sequencing according to the manufacturer’s instruction to identify the interacting protein partner(s). The *AD-T-antigen* and *BD-p53* interaction was considered as positive control.

### Bimolecular Fluorescence Complementation (BiFC) assay

The *Osr40c1* gene was cloned between the *StuI* and *BamHI* RE sites of *pVYCE (R)* vector to obtain *35S::Osr40c1-cVenus* construct. Simultaneously, the partners like Os*SAP8*, OsMNB1B, OsSAM2, and *OsH4* were cloned at the same RE sites of *pVYNE (R*) vector to generate *35S::OsSAP8-nVenus, 35S::OsMNB1B-nVenus, 35S::OsSAM2-nVenus*, and *35S::OsH4-nVenus* constructs respectively (Waadt et al. 2008). The recombinant constructs were transformed into *Agrobacterium* strain GV3101 and used for infiltration of onion epidermal cells in pairwise combinations according to Yang *et al*. (2014). The samples were analyzed to detect the expression of Venus protein under laser scanning confocal microscope (Olympus FV1000-IX81) using excitation wavelength of 514 nm. The *35S::Osr40c1-cVenus* construct with empty *35S:: nVenus* vector was considered as negative control.

### Homology modelling of *Os*r40c1, *Os*SAM2, *Os*SAP8, *Os*MNB1B and *Os*H4 proteins and molecular docking analysis

The protein sequences for *Os*r40c1, *Os*SAM2, *Os*SAP8, *Os*MNB1B, and *Os*H4 were retrieved from Uniprot and subjected to ROBETTA (Song et al., 2013) for prediction of 3D structures through homology modelling. The 3D structures were filtered or energy minimized by YASARA (Kreiger et al., 2009). Models were further validated by PROCHECK (Laskowski et al., 1996) using Ramachandran plot. According to Ramachandran plot, the residues of disordered region were refined using MOD Refiner and Sphinx servers (Fischer and Sali 2003; Dunbar et al., 2016) and further validated by PROCHECK. The molecular docking analysis for *Os*r40c1 and each of the interacting proteins were carried out by ClusPro software (Vazda et al., 2017; Kozakov et al., 2017).

### Virus Induced Gene Silencing (VIGS) of transgenic tobacco plants

Each fragment of *NtPDS, NtSAP8, NtSAM2, NtHMG1/2* genes were amplified from cDNA of tobacco using gene specific primers (supplementary table 1) and cloned into *pTRV2* vector between *EcoRI* and *BamHI* RE sites. The *Agrobacterium-mediated* inoculation of tobacco was carried out following Atsumi et al. (2018). For leaf infiltration, each *Agrobacterium* culture harbouring *pTRV1* and recombinant *pTRV2* constructs were mixed into 1:1 ratio and infiltrated into the leaves of transgenic tobacco (*Osr40c1* ectopic line) using a needleless syringe. The plants were maintained at 25 °C for effective viral infection and spread. The *NtPDS* gene was used as an internal control of VIGS.

### Statistical analysis

Three independent biological replicates and three technical replicates for each set were considered for all experiments where applicable and data were represented as mean ± standard error of mean (SEM). The differences in morphological parameters, metabolite contents and transcript abundances among genotypes and treatments were analyzed using GraphPad Prism version 8.0.0 software (GraphPad Software, San Diego, California USA) following two-way ANOVA and Sidak’s multiple comparison tests. The statistical significance at p ≤ 0.05 was considered to identify the difference between the two sets of data.

## Results

### *Osr40c1* expression is highly correlated with the degree of drought tolerance in rice

To analyze the drought tolerance potential of different rice cultivars, we have exposed eight *indica* rice cultivars to 7 days of drought stress and arranged them according to their degree of drought tolerance (Fig. 1A). Different agronomic traits like plant height, number of tillers, number of leaves, percentage of rolled leaves, percentage of brown leaves, root and shoot dry weight have been considered for this study (Fig. S1). As expected, no alteration in the plant height, number of tillers and leaf numbers have been observed in any of the cultivars. Leaf rolling and browning are considered as important physiological phenomena that determine the degree of drought stress impact on plants (Clarke, 1986). The leaf rolling has been observed after 48 hours of drought exposure in most of the varieties except IR36 and IR72. The percentage of rolled and brown leaves has been found to be the highest in MTU1010 and Jaldi13 and lowest in IR36 and IR72 in response to drought stress. Plant growth was markedly reduced after drought stress exposure in most of the varieties. The shoot and root dry weight differences had been found to be least in IR36 and IR72 and most prominent in MTU1010 and Jaldi13 in response to drought stress. After 24 hours of re-watering, the most pronounced revival has been observed for IR36 and IR72 as evidenced by the decreased percentage of rolled leaves and brown leaves. Next, we have estimated different metabolite contents from these eight cultivars in response to drought and re-watering. Significant augmentation in the contents of proline, glycine betaine, soluble sugar, ascorbic acid, and GSH has been observed in the roots and shoots of most of the cultivars in response to drought stress (Fig. S2). On the basis of all these analyses, the IR36 and IR72 cultivars have appeared to be most tolerant to drought stress, Jaldi13 and MTU1010 to be most susceptible while the other cultivars as moderately susceptible to drought stress.

**Fig. 1.**
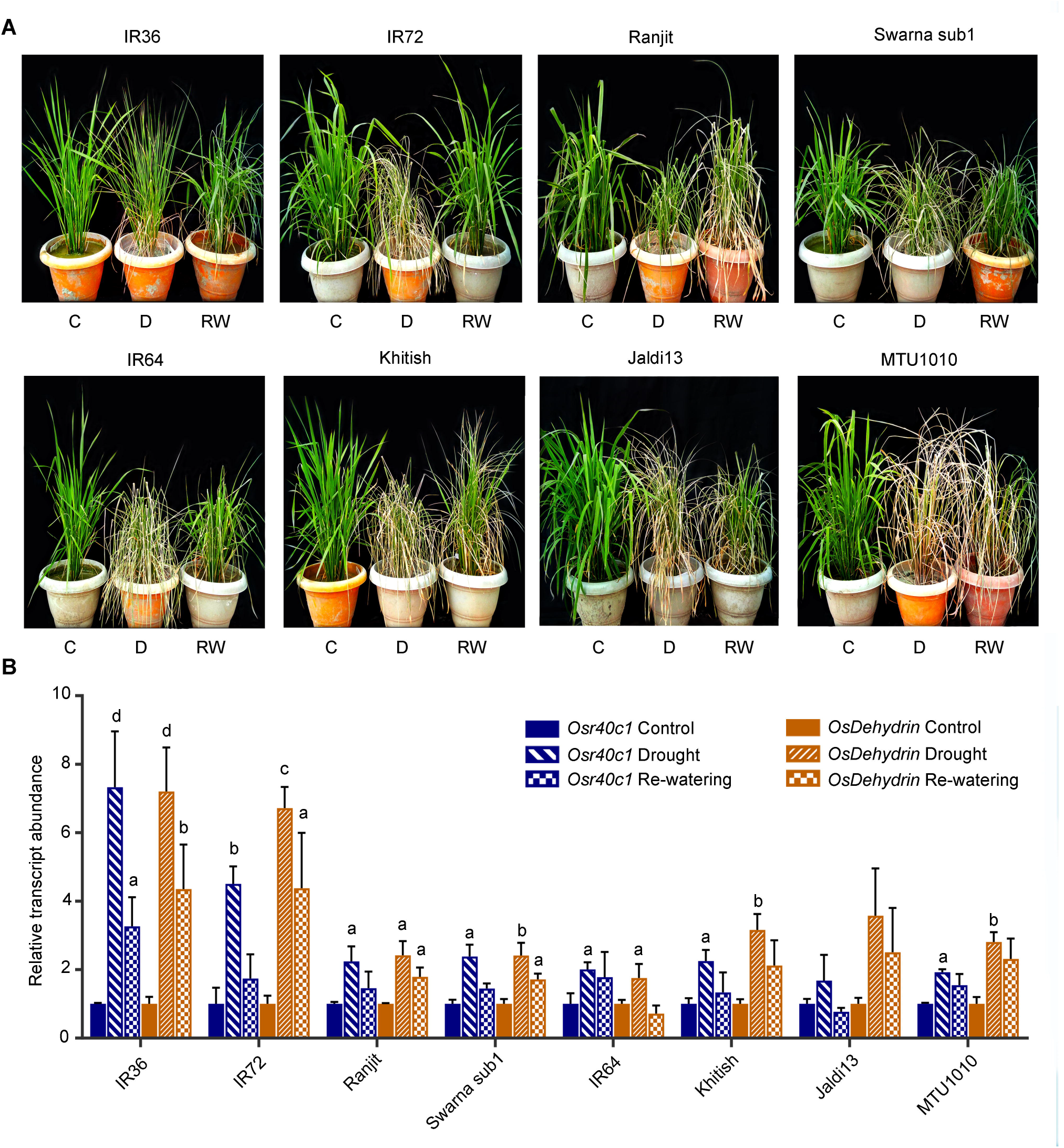
Drought stress analysis of eight *indica* rice cultivars, IR36, IR72, Ranjit, Swarna Sub-1, IR64, Khitish, Jaldi 13, MTU1010. 60 days old plants were exposed under drought stress for 7 days and different morphological parameters were analyzed. (A) Morphological responses of eight indica rice cultivars after 7 days of drought stress and 24 hrs of rewatering. (B) qRT-PCR analysis to study the relative transcript abundance for Osr40c1 protein and *Osdehydrin* genes. Results were represented as mean±SEM (n=3). Statistical difference between the cultivars under control and drought stress was denoted by small alphabet at p<0.05 (a), p<0.01 (b), p<0.001 (c) and p<0.0001 (d).

It has been observed that the *Osr40c1* expression is significantly higher in the root in comparison to the shoot under both control and drought conditions (Fig. S3). To determine its transcript abundance in the eight rice cultivars, we have performed qRT-PCR analysis for the *Osr40c1* gene as well as a marker gene, *OsDehydrin*, from the roots under control, drought and re-watered conditions (Fig. 1B). In response to drought stress, the highest induction in *Osr40c1* expression has been observed in case of IR36 (7.325 fold) and IR72 (4.505 fold) while the weakest induction has been noticed in case of MTU1010 (1.923 fold) and Jaldi13 (1.681 fold). In all other varieties, the induction level was moderate. In response to re-watering, the transcript abundance reduced to 3.62 fold in IR36 and 1.74 fold in IR72 suggesting their revival from stress. The expression of the marker gene, *OsDehydrin*, has also been found to be highest in the IR36 and IR72 as compared to the rest of the cultivars. These observations have interestingly demonstrated a positive correlation of *Osr40c1* expression with the degree of drought tolerance among the eight selected rice cultivars.

### *Osr40c1* gene is also regulated under PEG-mediated osmotic stress

To study if *Osr40c1* is regulated under osmotic stress as well, we have investigated the responses of the eight rice cultivars in response to 20% PEG (6000) exposure. The osmotic stress tolerance potential has been found to be highest in IR36 and IR72 and lowest in Jaldi13 and MTU1010 which supported our earlier observation (Fig. 2A). We have also measured the compatible solute and other metabolite contents from the eight cultivars. The alteration of metabolite contents in response to PEG mediated stress has been found to be similar to that of the drought stress (data not shown). We have next, analyzed the expression of the *Osr40c1* gene under control, PEG, and revival conditions. As expected, the transcript abundance was maximum in the case of IR36 (2.404 fold) and IR72 (2.724 fold) and minimum in the case of Jaldi13 (1.574 fold) and MTU1010 (1.540 fold) in response to PEG exposure (Fig. 2B). The transcript abundance dropped considerably in the tolerant cultivars but not in the susceptible ones in response to revival.

**Fig. 2.**
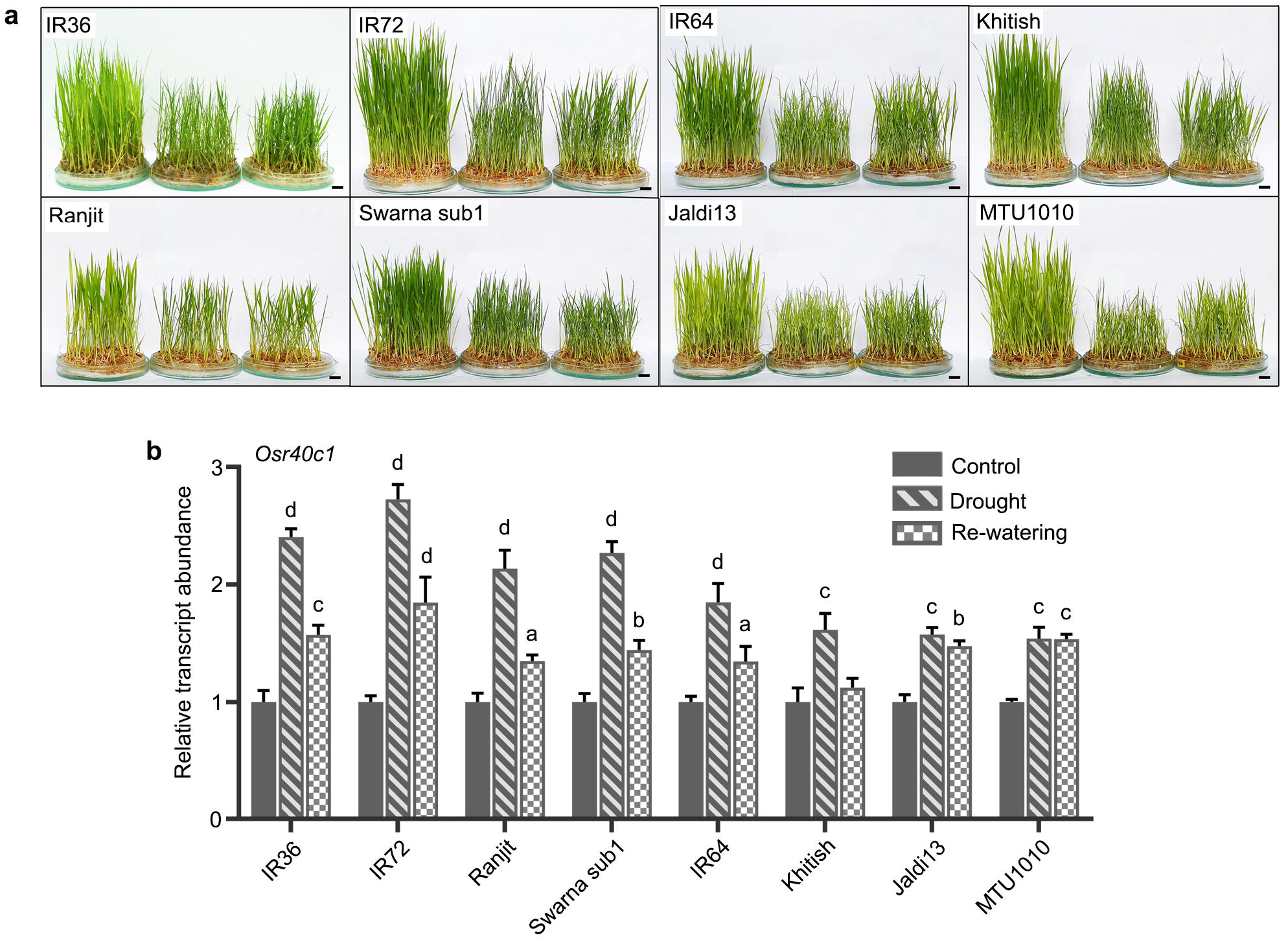
PEG mediated osmotic stress analysis of eight *indica* rice cultivars, IR36, IR72, Ranjit, Swarna Sub-1, IR64, Khitish, Jaldi 13, MTU1010. Two weeks of old plants were treated with PEG solution for 7 days. (A) Morphological responses of eight indica rice cultivars after 7 days of PEG treatment and 24 hrs of revival. (B) Samples were also used for qRT-PCR analysis to study the relative transcript abundance for *Osr40c1* and *Osdehydrin* genes. Results were represented as mean±SEM (n=3). Statistical difference between the cultivars under control and drought stress was denoted by small alphabet at p<0.05 (a), p<0.01 (b), p<0.001 (c) and p<0.0001 (d).

### *Os*r40c1 protein displays a nucleo-cytoplasmic localization

To determine the subcellular localization of the *Os*r40c1 protein, we have used the *35S::Osr40c1-eGFP* construct for *Agrobacterium*-mediated infiltration of onion epidermal cells. It has been observed that *Os*r40c1-eGFP fusion protein is predominantly localized in the nucleus and the cytoplasm (Fig. 3). A *35S::eGFP* construct has been used as a positive control. Onion cells, stained with DAPI confirmed the presence of the fusion protein in the nucleus.

**Fig. 3.**
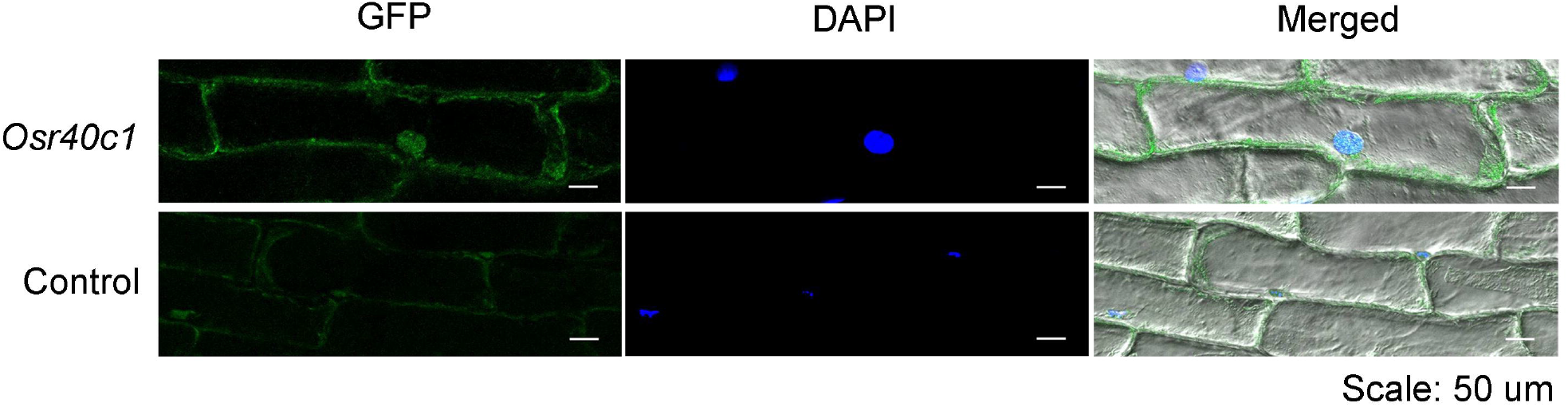
Localization of Osr40c1 protein in onion epidermal cells. Onion epidermal cells was infiltrated with recombinant construct (Osr40c1) and empty vector (control). Strong green fluorescence of GFP protein in case of recombinant construct indicates the nucleo cytoplasmic localization of r40c1 protein while in case of empty vector (pEGAD) very low green fluorescence is observed. Blue fluorescence of DAPI indicates nuclear localization.

### Overexpression of *Osr40c1* significantly enhances drought tolerance in rice

To biologically validate the drought-responsive role of *Osr40c1*, we have generated *Osr40c1* overexpressing transgenic rice lines by introducing the *35S::Osr40c1-eGFP* construct via *Agrobacterium-mediated* transformation. Out of 17 independent putative T0 transformed plants, 13 were found to be positive (Fig. S4). Three best lines have been selected based on the expression level of *Osr40c1* and maintained up to T2 generation. All the selected overexpression (OX) lines have been found to exhibit significantly improved drought stress tolerance over the wild-type (WT) and vector control (VC) plants (Fig. 4A). We have analyzed different agronomical parameters like plant height, tiller numbers, percentage of brown and rolled leaves, shoot and root dry weight of the OX lines and compared with the WT and VC plants. The OX lines displayed no morphological alteration as compared to the WT under control condition. However, under drought stress, the OX lines have been found to display a lower percentage of leaf rolling and brown leaves as compared to the VC and WT plants (Fig. S5). Furthermore, we have measured the proline, glycine betaine, soluble sugars, ascorbic acid, and GSH contents from the roots and shoots of the OX, VC, and WT plants. The metabolite accumulation has been found to increase in all the lines in response to drought stress (Fig. S6).

**Fig. 4.**
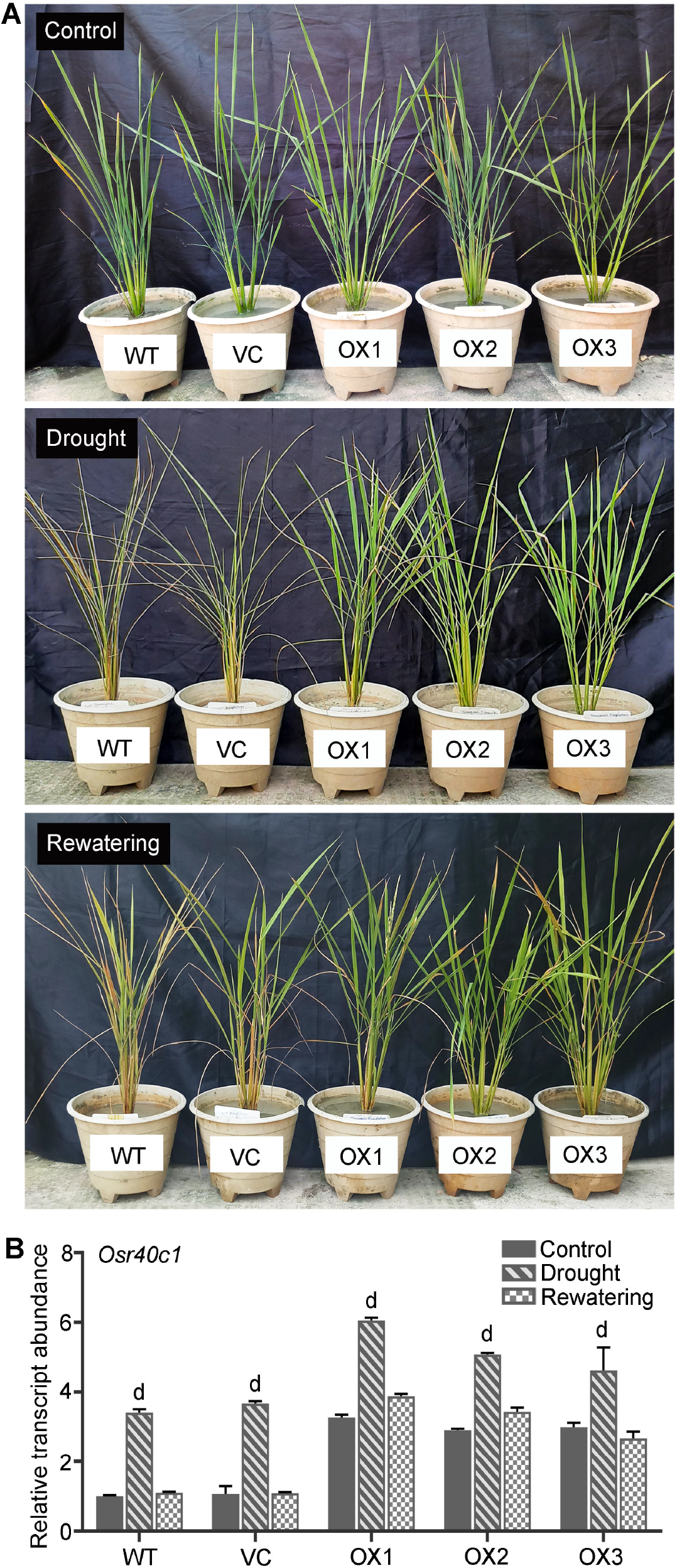
Analysis of transgenic rice plants overexpressing *Osr40c1* gene. The WT, VC and three independent transgenic lines (OX1, OX2 and OX3) were subjected to drought stress for 7 days and (A) morphological responses was recorded. The relative transcript abundance of *Osr40c1* gene (E) was also analyzed. Results were represented as mean±SEM (n=3). Statistical difference between the lines under control and drought stress was denoted by small alphabet p<0.0001 (d).

Next, the expression of *Osr40c1* has been analyzed under control, drought and re-watered conditions. The expression of the *Osr40c1* gene has been found to be increased by around 3 fold in the OX lines as compared to the WT under control condition. Under drought stress as well, the transcript abundance has been found to be significantly higher in the OX lines over the VC and WT plants (Fig. 4B).

### Ectopic expression of *Os*r40c1 enhances drought tolerance in tobacco

Further, to confirm the function of *Os*r40c1 in a heterologous system, tobacco lines ectopically expressing the *Osr40c1* gene have been developed. Out of 21 putative transformed lines generated, 16 positive transformed lines have been obtained (Fig. S7). Three best lines with the highest *Osr40c1* expression have been selected for further analyses. The transgenic tobacco (OX) lines have displayed significant improvement in drought tolerance over the WT and VC lines when exposed under drought stress (Fig. 5A). Furthermore, a significant accumulation of *Osr40c1* transcript has been observed in the transgenic lines while no expression could be detected in the WT and VC plants since *r40c1* is exclusively present in rice (Fig. 5B). These observations, together, have strongly suggested that *Os*r40c1 plays a crucial role in imparting drought stress tolerance in plants.

**Fig. 5.**
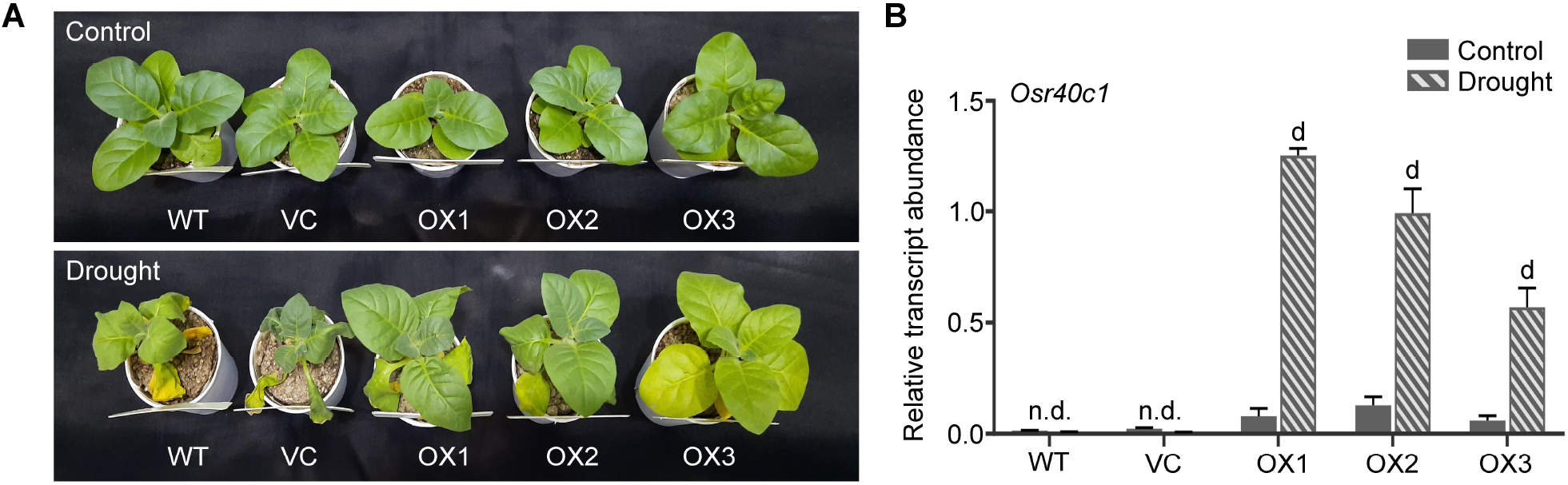
Analysis of transgenic tobacco plants. The WT, VC and three independent transgenic lines (OX1, OX2 and OX3) were subjected to drought stress for 5 days and (A) morphological responses was recorded. The relative transcript abundance of *Osr40c1* gene (E) was also analyzed. Results were represented as mean±SEM (n=3). Statistical difference between the lines under control and drought stress was denoted by small alphabet p<0.0001 (d).

### *O*sr40c1 interacts with several drought-responsive protein partners

To identify the interacting partners of *Os*r40c1 protein, we have performed yeast two-hybrid analysis using *Os*r40c1 protein as bait. The cDNA library prepared from rice roots under drought stress has been used as prey. After performing the yeast two-hybrid analysis and screening on DDO, QDO, and high stringency QDO/X/A plates, 16 blue colonies were selected. Sequence analysis identified 8 non-redundant protein partners for *Os*r40c1 namely, *Os*MNB1B (LOC4342129), *Os*SAP8 (LOC4341520), *Os*SAM2 (LOC4326996), OsH4 (LOC4347135), uncharacterized-I (LOC4337962), uncharacterized-II (L0C4338275), *Os*PBL19 (LOC4342066), and *Os*CyclinD (LOC4331985) (Fig. 6A). The interaction of the p53 protein with T-antigen has been used as a positive control. Additionally, we have checked the expression levels of each of these partners along with *Osr40c1* from rice roots in response to drought stress. Fascinatingly, it has been observed that the expression of *Osr40c1, OsMNB1B, OsSAM2* and *OsH4* are significantly up-regulated under drought stress thus indicating their possible involvement in regulating drought response in plants (Fig. 6B).

**Fig. 6.**
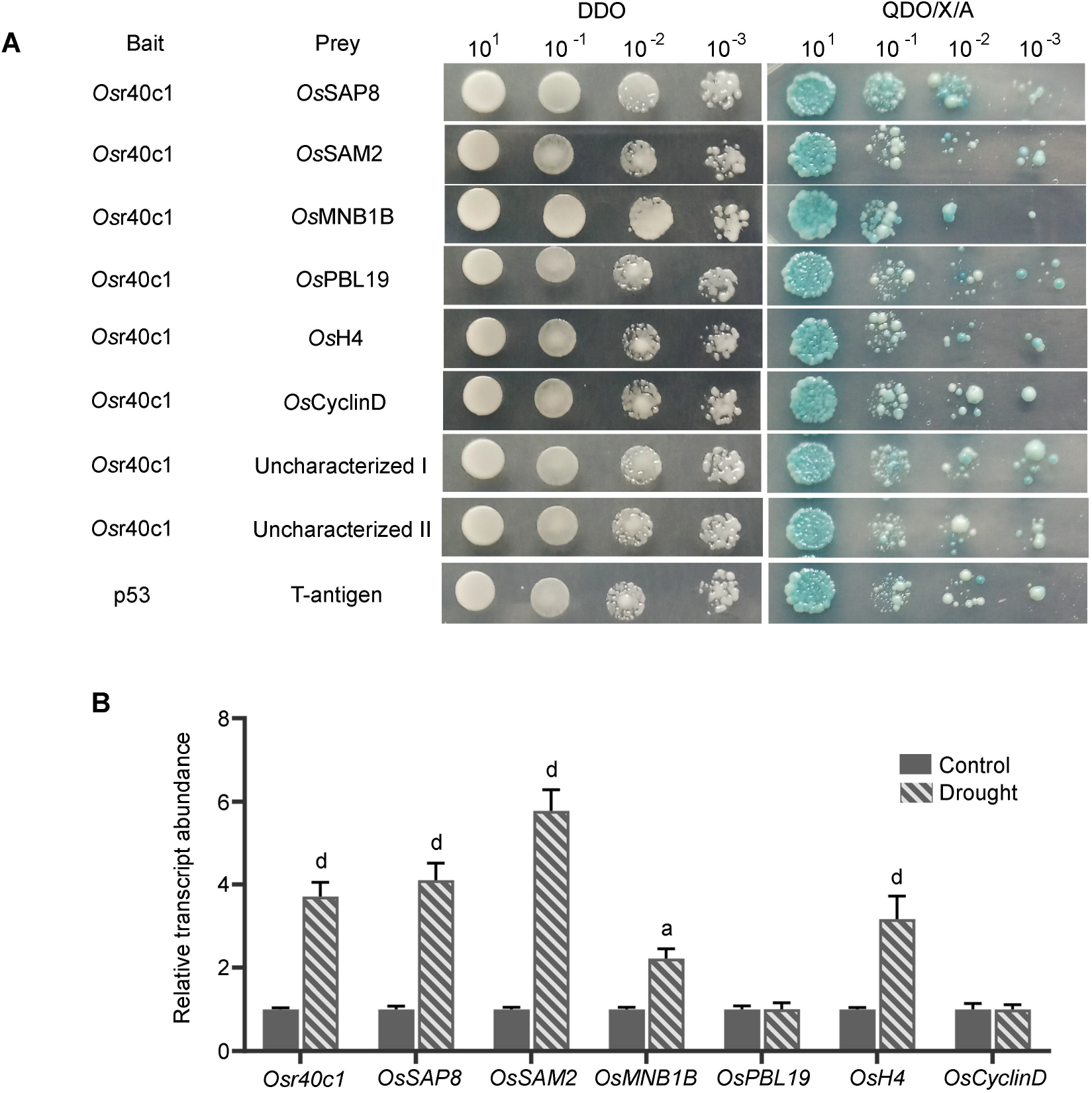
Identification of interacting protein partners of *Os*r40c1 protein under drought stress. (A) Yeast two-hybrid analysis identified the interaction of *Os*r40c1 with eight different proteins like OsSAP8, OsSAM2, OsMNB1B, OsPBL19, OsH4, OsCyclinD, Uncharacterised I and Uncharacterised II proteins. The interaction of p53 protein with T-antigen was used as a positive control. (B) qRT-PCR analysis to study the relative transcript abundance of each interacting protein partners along with *Osr40c1* under drought stress. Results were represented as mean±SEM (n=3). Statistical difference between the cultivars under control and drought stress was denoted by small alphabet at p<0.05 (a) and p<0.0001 (d).

To validate the interaction of these 4 selected protein partners, we have performed BiFC analysis in onion epidermal cells. A strong yellow fluorescent signal of Venus protein has been observed in the cytoplasm as well as the nucleus when *Os*SAP8 interacted with *Os*r40c1 protein. On the other hand, the interaction of *Os*SAM2, *Os*MNB1B, and *Os*H4 with *Os*r40c1 has predominantly been detected in the nucleus (Fig. 7). No fluorescent signal has been observed for the empty *35S::nVenus* vector and *35S::Osr40c1-cVenus* pair which served as the negative control.

**Fig. 7.**
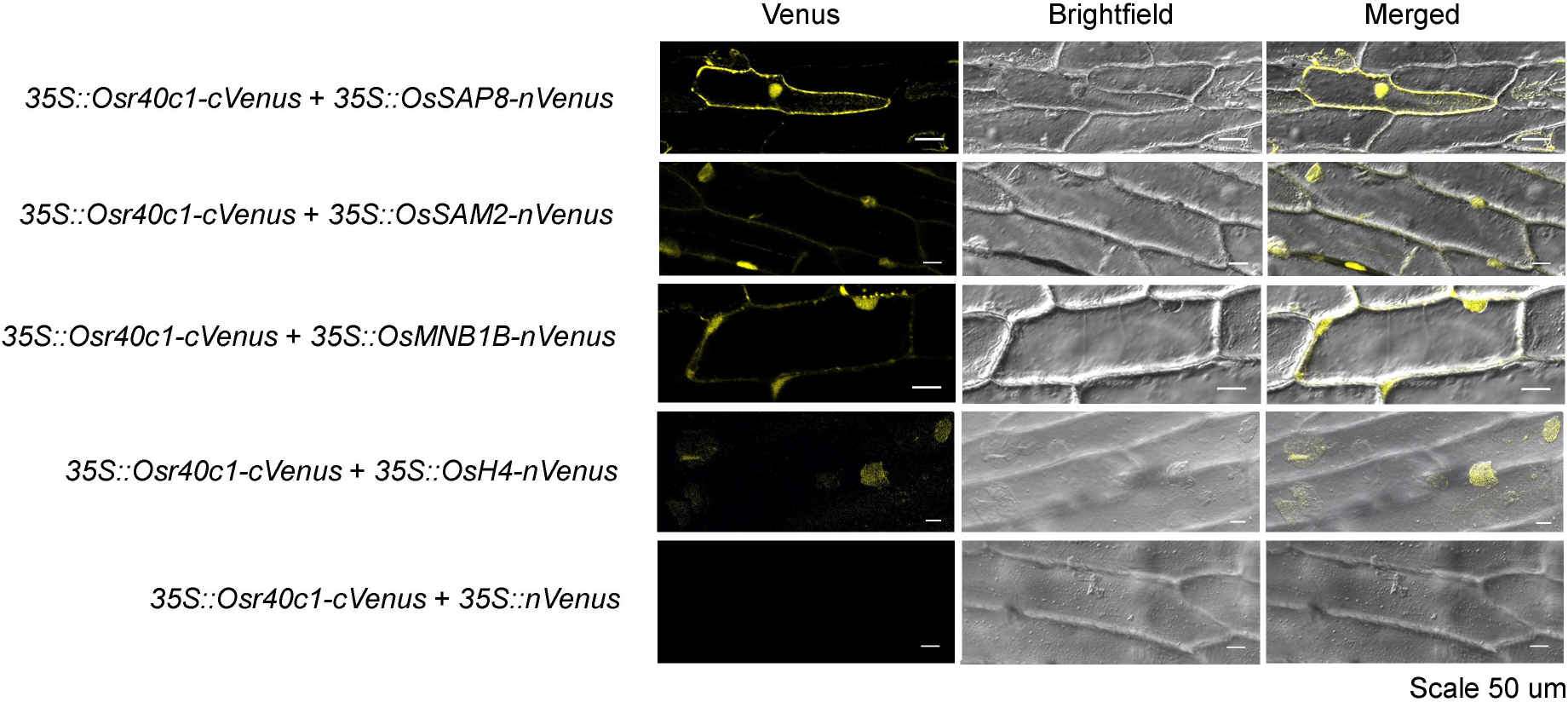
BiFC analysis of Osr40c1 and interacting protein partners. The interaction of *Os*r40c1 protein with *Os*SAP8 in the nucleus and cytoplasm and with *Os*SAM2, OsMNB1B and OsH4 in the nucleus was found. Venus fluorescence, bright field, and merged images were represented for each set of constructs.

*In-silico* analysis has been performed through homology modeling followed by molecular docking analysis to re-confirm the interaction of the protein partners with *Os*r40c1. The 3D structures for *Os*r40c1, *Os*SAP8, *Os*H4, *Os*SAM2, and *Os*MNB1B have been generated using Robetta server followed by structure refining, energy minimization and validation using PROCHECK server (Fig. S8). The percentage of amino acid residues that fall in the most favored regions of the Ramachandran plot has been found to be 88%, 84.9%, 94%, 88.7% and 94.2% respectively, while none of the residues have been found in the generously allowed or the disallowed regions (Fig. S9-S13). The best 3D structures for each protein have been selected and used for molecular docking analysis which re-confirmed the interaction of *Os*r40c1 with each of its interacting protein partners (Fig. 8).

**Fig. 8.**
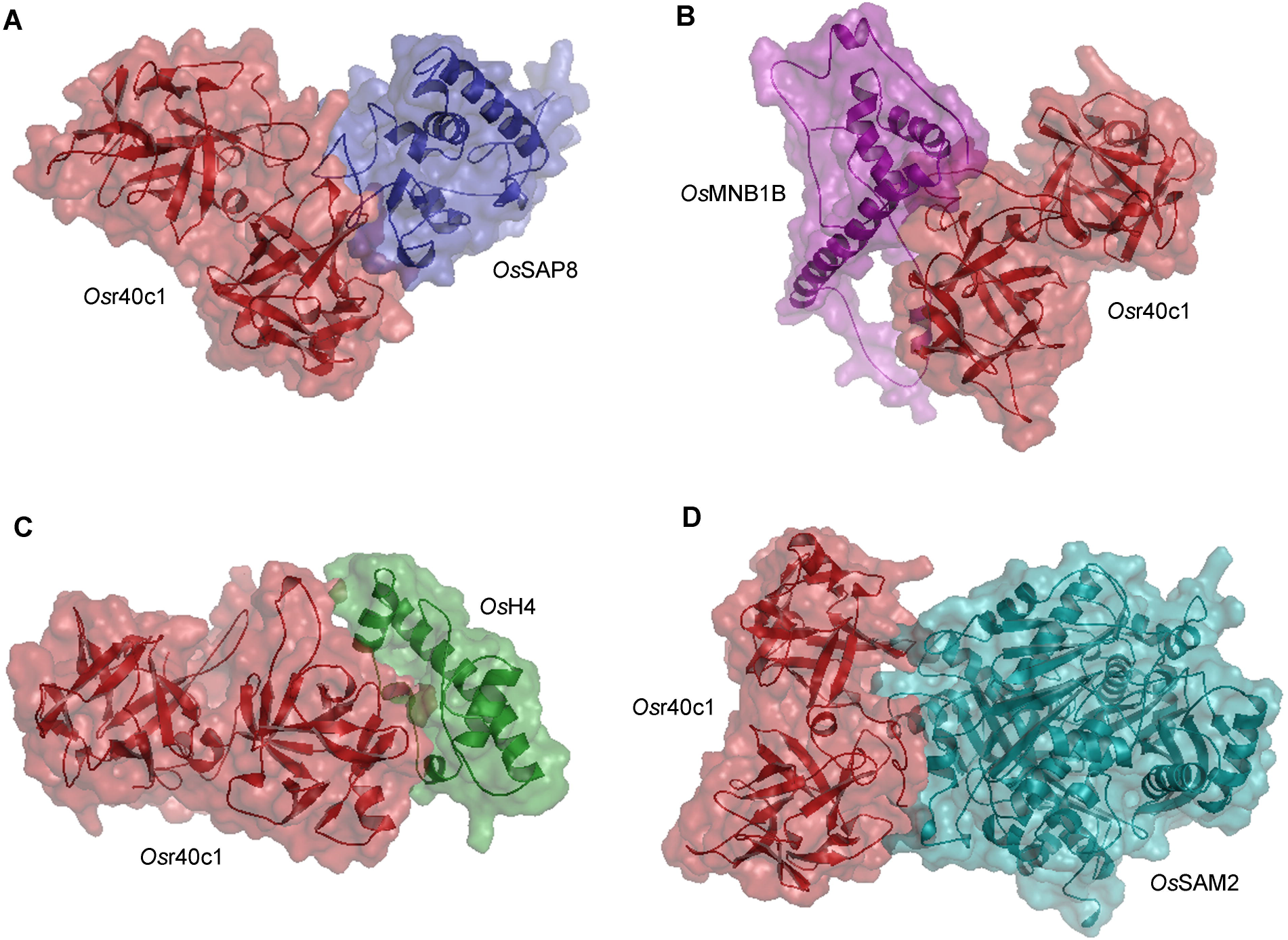
*In silico* analysis for *Os*r40c1-OsSAP8, Osr40c1-OsSAM2, Osr40c1-OsMNB1B and Osr40c1-OsH4 interaction. Protein structures for *Os*r40c1, OsSAP8, OsSAM2, OsMNB1B, and OsH4 were generated through homology modelling. The structures were used for molecular docking analysis which confirms the interaction.

### Silencing of the interacting protein partners leads to drought susceptibility in transgenic tobacco plants

To unravel the biological significance of the protein partners in *Os*r40c1-mediated drought response in plants, the orthologs of *Os*SAP8, *Os*SAM2 and *Os*MNB1B have been silenced through VIGS in the transgenic tobacco lines ectopically expressing *Osr40c1*. The proteins *Nt*HMG1/2 (LOC107781188), *Nt*SAP8 (LOC107822754) and *Nt*SAM2 (LOC107770931), have been considered as orthologs to the rice proteins *Os*MNB1B, *Os*SAP8, and *Os*SAM2 respectively. The silencing of *NtPDS* has been used as a positive control for VIGS system (Ratcliff et al., 2001; Turnage et al., 2002). After 7 days of drought treatment, the VIGS lines for *NtHMG1/2, NtSAP8* and *NtSAM2* have displayed prominent drought sensitivity in the otherwise tolerant transgenic lines (Fig. 9A,C,E,G). Severe wilting has been noticed in the case of all three silencing events under drought stress. Interestingly, plants of *NtSAP8* and *NtHMG1/2* silencing lines have been found to be more affected than WT under drought stress. The relative expressions of the silenced genes (*NtHMG1/2, NtSAP8* and *NtSAM2*) along with *Osr40c1* gene in all three lines have been analyzed. A reduction by 82.71%, 87.739%, and 89.83% has been observed in case of *NtHMG1/2, NtSAP8 and NtSAM2* transcript abundance respectively in the silenced plants. However, significant induction of *Osr40c1* has been observed in the silenced as well as the non-silenced lines under drought stress (Fig. 9B,D,F). Together, our analysis strongly indicates that interaction with these protein factors is crucial for the *Os*r40c1-mediated stress tolerance *in planta*.

**Fig. 9.**
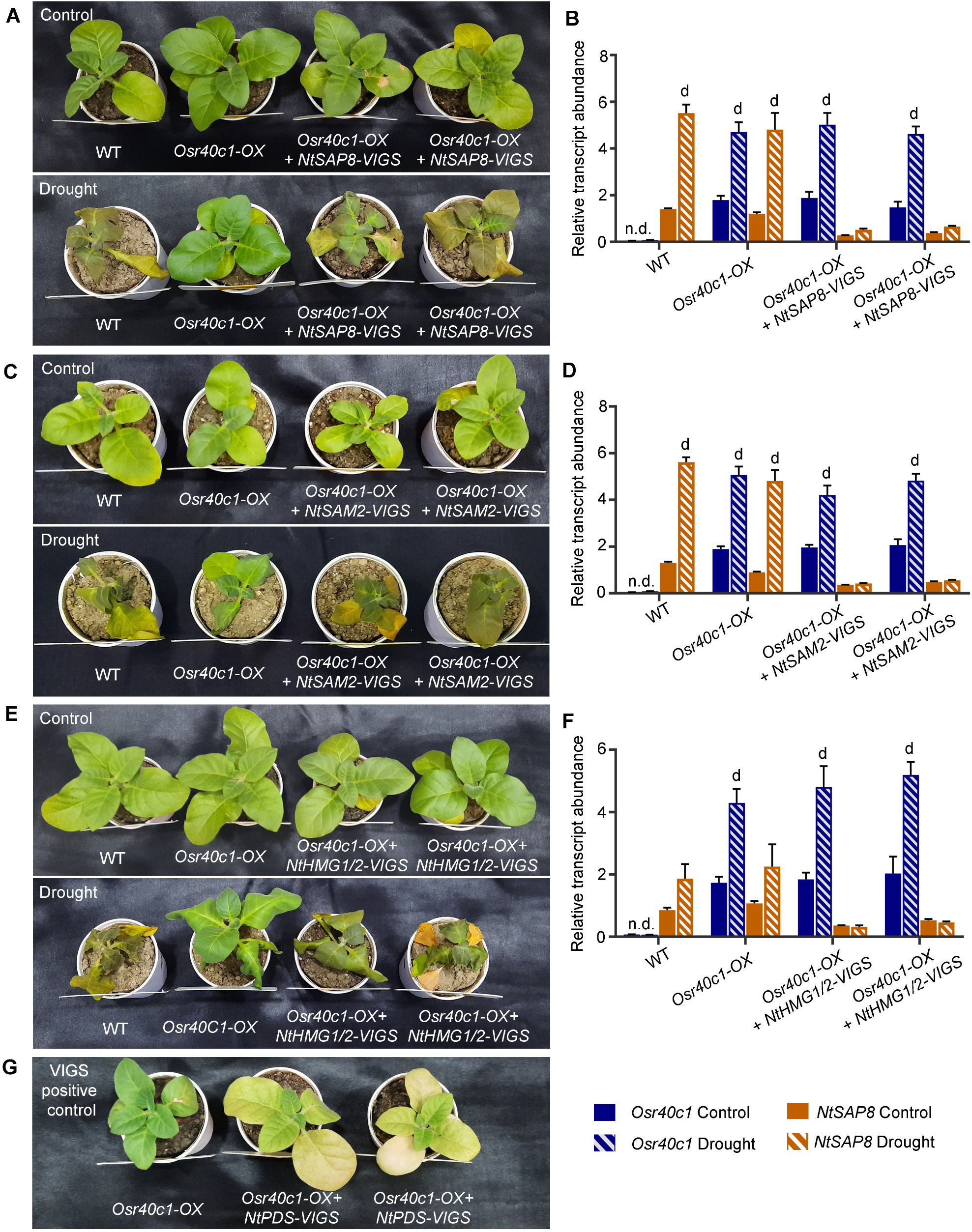
Analysis of tobacco ectopic lines after VIGS mediated silencing of interacting protein partners. Morphological responses of NtSAP8 silenced lines (A) NtSAM2 silenced lines (C) and (E) NtHMG1/2 silenced lines after 5 days of drought exposure. qRT-PCR analysis to study the relative transcript abundance of NtSAP8 (B), NtSAM2 (D) and NtHMG1/2 (F) gene along with *Osr40c1* under drought stress. Results were represented as mean±SEM (n=3). Statistical difference between the cultivars under control and drought stress was denoted by small alphabet at p<0.0001 (d).

## Discussion

Lectins are a family of carbohydrate-binding proteins known to play vital roles in diverse physiological phenomena in plants. Among them, the R40 proteins are considered as a stress-responsive group of lectin proteins that serve predominant roles in regulating osmotic stress responses in different plant species (Moons *et al*., 1997; Jiang *et al*., 2012). Drought is a major threat for agriculture and hampers rice productivity. In plants, roots are the first to respond to water deficiency in soil and triggers downstream signalling pathways for its adaptation under drought stress. Excitingly, the *Os*r40c1 is predominantly expressed in the roots under control as well as drought conditions. This observation also corroborates with the earlier report which identified the up-accumulation of *Os*r40c1 protein in rice roots under drought stress (Paul *et al*., 2015). However, the mechanism of how this protein imparts drought stress tolerance in plants remains elusive so far.

In the present study, we have demonstrated that the expression of *Osr40c1* gene is highly correlated with the degree of drought tolerance in rice. We have studied 8 *indica* rice cultivars for their drought and osmotic stress tolerance potentials. Several morphological parameters have been analyzed under control, stress and re-watered conditions. Among them, the percentage of rolled and brown leaves, increase in root length and changes in the root and shoot biomass in response to stress has been considered to assess their drought tolerance potential. Out of the 8 varieties, the IR36 and IR72 considered as drought tolerant owing to their lower percentage of rolled and brown leaves, higher biomass, and longer root length in response to drought stress. The two varieties, Jaldi13 and MTU1010 have displayed highest degree of drought susceptibility. The accumulation patterns of different osmotically active metabolites have also been analyzed in these varieties. Proline and glycine betaine are considered as osmotically active compounds that maintains cellular integrity during osmotic stress responses in plants (Chen *et al*., 2002). The involvement of GSH and ascorbate in regulating abiotic stress has also been widely reported (Noctor and Foyer, 1998; Datta et al., 2020). The highest accumulation of these compounds in IR36 and IR72 suggest their drought tolerant properties. However, in the case of all the 8 cultivars, the accumulation of these metabolites has increased in response to drought stress. The expression of the *Osr40c1* gene has been observed to increase under drought stress condition in all the cultivars. Interestingly, its transcript abundance has been the highest in the tolerant cultivars, IR36 and IR72 and the lowest in the susceptible cultivars, Jaldi13 and MTU1010 thus indicating a positive correlation with the drought tolerance potential in plants. Moreover, the transgenic rice lines overexpressing *Osr40c1* gene exhibited significantly improved drought tolerance over the WT and VC plants further establishing its role in drought response. A similar observation has been found in case of tobacco plants ectopically expressing the *Osr40c1* gene.

Once the crucial role of *Os*r40c1 has been established in drought tolerance, we have aimed to unravel the mechanism of how it imparts this tolerance in plants. Since protein localization may have important influence on protein function, we have first analyzed the sub-cellular localization of this protein. Fascinatingly, the protein has been found to exhibit a nucleo-cytoplasmic localization. Next, several interesting protein partners have been identified to interact with the *Os*r40c1 protein. In addition, two uncharacterized proteins have also been identified which needs to be explored in future. Out of the 8 identified partners, a transcription factor, *Os*SAP8, and 2 chromatin-associated proteins, *Os*MNB1B and *Os*H4 have been found to be drought-responsive. Silencing of each these partner proteins have resulted in pronounced drought susceptibility of the transgenic tobacco plants ectopically expressing *Osr40c1* gene. This observation confirms that these protein partners are crucial for the *Os*r40c1-mediated drought stress tolerance in plants.

*Os*SAP8 is an osmotic stress-responsive transcription factor that comprises of two DNA-binding zinc finger domains – an N-terminal AN20 domain and a C-terminal AN1 domain. Previous studies have reported that several SAP proteins including SAP8 are regulated by drought and salinity stress and they enhance drought tolerance in rice (Kanneganti and Gupta 2007; Kothari *et al*., 2016). It has also been reported that ABA triggers the accumulation of *Os*SAP8 in plants. Therefore, it can be hypothesized that the *Osr40c1*, being an ABA responsive gene as well, can regulate *Os*SAP8 via a common ABA signalling pathway. Moreover, it has been demonstrated that the A20 domain of SAP proteins can interact with itself while the AN1 domain can interact with the A20 domain (Kanneganti and Gupta 2007). In addition, the *Os*SAP8 imparts drought stress tolerance in plants via interaction with other stress associated proteins to regulate the intricate signalling mechanism of drought tolerance (Giri *et al*. 2011). Therefore, it can be assumed that the *Os*r40c1 interaction leads to a conformational change in the *Os*SAP8 protein thus facilitating its binding to the *cis*-acting regulatory regions of drought responsive genes.

The *Os*r40c1 protein has also been found to interact with a chromatin-associated protein *Os*MNB1B, which is known to participate in chromatin modification and transcriptional induction. The *Os*MNB1B protein belongs to the high mobility group (HMG) proteins and bears a high sequence homology with the HMGB1 protein. It has been widely demonstrated that the HMGB1 protein can compete with the histone H1 to bind to the linker nucleosomal region of DNA. This opens up the chromatin and makes the DNA accessible to transcription factors (Nightingale *et al*., 1996; Catez *et al*., 2004). Earlier, it has been reported that maize MNB1B can bind to a specific AAGG motif in DNA whereas rice HMG1B protein specifically interacts with the four way regions (4H) of DNA and DNA minicircle thus leading to bending of the DNA molecule (Yinagisawa and Izui 1993; Wu *et al*., 2003). Besides, the HMG1 protein has been found to be interacting with the bZIP transcription factors and to enhance their binding to their target regulatory elements in DNA (Izawa et al., 1994). In addition, the HMGB1 proteins have been reported to function in an ABA-responsive pathway to impart abiotic stress tolerance (Christov *et al*., 2007). Keeping in view all these interesting findings, it can be attributed that the *Os*MNB1B protein helps in chromatin remodelling of drought responsive genes thus making them accessible to the *Os*SAP8-Osr40c1 complex which ultimately leads to the induction of the downstream drought responsive genes. In addition, this entire pathway may be operated via an ABA-responsive pathway. The histone protein modification has been commonly associated with several osmotic stress responsive genes in various plant species (Kim *et al*., 2015). This modification includes methylation and acetylation of different lysine and arginine residues of histone H3 and H4. Since *Os*H4 has been identified as one of the interacting partners of *Os*r40c1, the probability of *Osr*40c1-mediated histone modification of different downstream drought-responsive genes cannot be ruled out.

SAM2 is an S-adenosine methyltransferase that is known to be up-regulated under drought stress and catalyze the synthesis of S-adenosyl-methionine (S-AdoMet). S-AdoMet is an important component of DNA, RNA, and protein methylation in plants (Meng *et al*., 2018). On the other hand, SAM2 helps in polyamine biosynthesis which is essential for plants to cope with the adverse conditions of drought or salt stress (Ma *et al*., 2017). The overexpression of sugar beet SAM2 also exhibited an enhanced drought tolerance in *Arabidopsis* (Ma *et al*. 2017). Together, it can be hypothesized that the *Os*r40c1 protein may activate *Os*SAM2 which leads to synthesis of more S-AdoMet. This in turn increases the polyamine biosynthesis ultimately leading to drought stress tolerance in plants.

In summary, it can be concluded that the rice lectin protein, *Os*r40c1, provides drought stress tolerance in rice by interacting with several exciting protein partners. It has been hypothesized that drought stress induces the expression of *Os*r40c1 protein. This protein then interacts with the chromatin-associated proteins, *Os*MNB1B and *Os*H4 presumably to induce chromatin remodelling. This enables the *Os*SAP8 transcription factor to bind to its target DNA motif to induce the expression of downstream drought-responsive genes (Fig. 10). Since all these proteins have been reported to be ABA-responsive, the entire pathway may function as a complex in an ABA-dependent pathway. In addition, *Os*r40c1 also interacts with *Os*SAM2 protein thus inducing drought tolerance via polyamine biosynthesis pathway. Together, the present investigation demonstrates the novel role of *Os*r40c1 in imparting drought tolerance via regulation of crucial transcriptional regulators as well as SAM2 protein in plants.

**Fig. 10.**
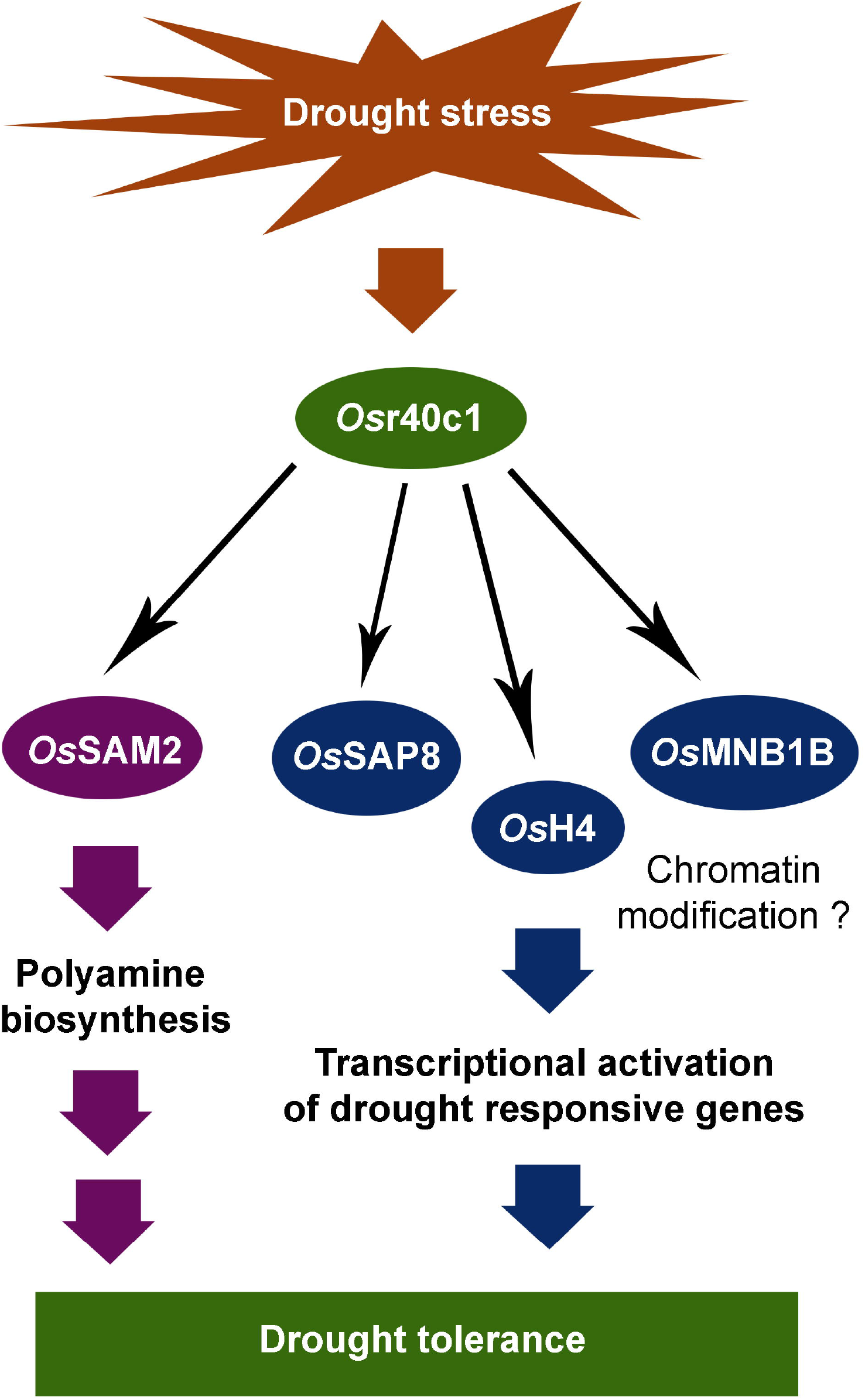
Model for *Os*r40c1-mediated regulation of drought stress. Drought stress triggers *Osr40c1* gene expression in cells. In the mean tine, the expression of *OsSAM2, OsSAP8, OsMNB1B* and *OsH4* genes are also induced followed by higher accumulation. The *Osr40c1* then binds with the chromatin modification associated proteins like OsSAP8, OsMNB1B and OsH4 to activate the transcription of downstream drought responsive genes presumably inducing the chromatin remodelling of drought responsive genes. On the other hand, Osr40c1 can interact with OsSAM2 to activate the protein that enhances the polyamine biosynthesis. Together, these results suggest both the transcriptional activation of downstream drought-responsive genes and polyamine accumulation which ultimately leads to drought stress tolerance in plants

## Supplementary data

Fig. S1. Morphological analysis of 8 *indica* rice cultivars in response to drought stress

Fig. S2. Estimation of different metabolites from 8 *indica* rice cultivars in response to drought stress

Fig. S3. Expression of *Osr40c1* gene from root and shoot of rice plant in response to drought stress

Fig. S4. Screening of transgenic rice lines overexpressing *Osr40c1* gene

Fig. S5. Morphological analysis of transgenic rice plants in response to drought stress

Fig. S6. Estimation of different metabolites from transgenic rice plants in response to drought stress

Fig. S7. Screening of transgenic tobacco lines ectopically expressing *Osr40c1* gene

Fig. S8. Homology modelling of *Os*r40c1, *Os*SAP8, *Os*SAM2, *Os*MNB1B, and *Os*H4 proteins

Fig. S9. Ramachandran plot analysis for *Os*r40c1 protein

Fig. S10. Ramachandran plot analysis for *Os*SAP8 protein

Fig. S11. Ramachandran plot analysis for *Os*H4 protein

Fig. S12. Ramachandran plot analysis for *Os*SAM2 protein

Fig. S13. Ramachandran plot analysis for *Os*MNB1B protein

Table S1. List of primers used

## Acknowledgement

This work has been supported by the Department of Science and Technology and Biotechnology, Government of West Bengal, India [BT(Budget)/RD-29/2016] and University Grant Commission, Government of India, India [UGC-CAS (Phase VII)]. We thank the central instrumentation facility of the Department of Botany, University of Calcutta and Dr. A. P. J. Abdul Kalam Government College as well as the confocal microscopic facility of DBT-IPLS, Department of Biochemistry, University of Calcutta. We thank Prof. Jörg Kudla (University of Munster, Germany) for providing the *pVYNE* and *pVYCE* vectors and Prof. Prabod Trivedi (CSIR-National Botanical Research Institute, India) for kindly sharing the *Agrobacterium tumefaciens* GV3101 strain. We are also thankful to Dr. Jyothilakshmi Vaddassery, Staff Scientist IV, NIPGR, New Delhi for providing us the *pTRV1* and *pTRV2* vectors.

## Author Contributions

RD and SP conceived and designed the research plan; SH performed most of the experiments; CR performed rice transformation experiment, RD performed the *in silico* analyses; RD and SP analyzed the data; SH drafted the manuscript; RD and SP supervised and complemented the writing.

